# Local Thermal Adaptation occurs via Modulation of Mitochondrial Activity in Chinook Salmon

**DOI:** 10.1101/2023.09.28.560008

**Authors:** Bradford Dimos, Alex Lopez, Patricia Schulte, Michael Phelps

## Abstract

Freshwater and anadromous fish have been identified as one of the most at-risk groups threatened by climate induced warming given their metabolic dependence on environmental temperature and limited ability to track favorable environmental conditions. The future of these fish will depend on their ability to adapt to new thermal regimes over biologically relevant timescales. However, the mechanistic understanding of temperature responses required to predict if adaption can keep pace with climate change is limited. To address this question, we investigated the mechanistic basis of thermal adaptation across multiple levels of biological organization in the iconic and endangered Chinook Salmon (*Oncorhynchus tshawytscha*). We uncovered a mechanistic basis of thermal adaptation centered on the thermal responsiveness of mitochondrial function which modulates the extent to which rising temperatures increase metabolic demand on the cardio-respiratory system. These insights demonstrate that the populations studied here are able to maintain high levels of physiological performance at temperatures several degrees above historic averages, indicating that the decline Chinook Salmon and failure to recover despite conservation efforts is unlikely to be due to increased temperatures as a consequence of climate change.

## Intro

Climate change threatens ecosystem stability by altering current thermal regimes, creating mismatches relative to the historical conditions to which many species have adapted (Ainsworth et al., 2016; Wernberg et al., 2016). These climate-change induced mismatches have the potential to worsen as the pace of climate change accelerates and the projected rise in temperatures outpaces the rate at which organisms can adapt. Aquatic ectotherms are particularly sensitive to rising temperatures given the dependence of metabolic rate on environmental temperature (Deutsch et al., 2015), with some predictions indicating that over half of all fish species will experience temperatures exceeding their thermal tolerance limits by 2100 (Dahlke et al., 2020). The vulnerability of specific species or populations is often assessed via oxygen budgets as the ability to meet the increased oxygen demand imposed by higher metabolic rates has been suggested to constrain thermal performance (Pörtner et al., 2017). Given this, mitochondrial activity and cardiorespiratory capacity have received significant interest from the research community for their role in influencing thermal performance. However, studies linking these across multiple levels of biological organization are lacking (Schulte, 2015) limiting the mechanistic understanding required to predict how organisms will respond to climate change (Urban et al., 2016).

Pacific salmon (genus, *Oncorhynchus*) are cold water species that are highly vulnerable to the negative impacts of climate change (Weiskopf et al., 2020). These iconic fish are essential to the culture, food security and economy of the North Pacific, and are threatened across their range, with rising stream temperatures highlighted as a major contributor to this decline (Crozier et al., 2021). The vulnerability of Pacific salmon to high temperature stems from their anadromous life history, which promotes local adaption of reproductively isolated populations to historical streams conditions as a result of their return to natal streams to spawn (Eliason et al., 2011; Farrell et al., 2008; Zillig, FitzGerald, et al., 2023). As cold-water species, even modest increases in temperature can have severe consequences on population viability by reducing cardio-respiratory performance, which has been widely suggested to constrain thermal tolerance in salmonids (Clark et al., 2008) (Eliason et al., 2013), however the mechanistic basis of this vulnerability is an open question. Chinook Salmon (*O. tshawytscha*) are the largest of the Pacific salmon, and are highly imperiled (Hoekstra et al., 2007), with several populations being listed on the United States Endangered Species Act and the Canadian Species at Risk Act. Thus, understanding the mechanisms underlying local thermal adaptation in Chinook is essential for understand the risk climate change poses to this species (Crozier et al., 2007).

To assess the vulnerability of *O. tshawytscha* to rising stream temperatures we sought to uncover the mechanistic basis of the locally adapted thermal performance displayed by salmon populations. Given the interdependence between temperature, metabolic rate and the cardiorespiratory system we hypothesized that differences in mitochondrial function may be a key axis of adaption in *O. tshawytscha* originating from warm versus cool streams. To address this hypothesis, we tested four populations originating from disparate thermal regimes for differences in aerobic performance, acute thermal tolerance, mitochondrial physiology, gene expression and genetic differentiation. Our investigation reveals that aerobic performance and acute thermal tolerance are linked, and that the expression of genes involved in ventricular contraction and mitochondrial function are associated with acute thermal tolerance and targets of selection. Finally, we demonstrate that the thermal responsiveness of mitochondria is reduced in warm-adapted populations and that this represents a key axis of thermal adaptation in *O. tshawytscha*. These results provide unparalleled mechanistic insight spanning multiple levels of biological organization into the development of locally adapted phenotypes in an endangered species. These results suggest that *O. tshawytscha* parr able to tolerate temperatures several degrees above what they currently experience suggesting that the observed declines and failure to recover populations are likely not due to an inability of salmon fry to handle rising temperatures, at least in the central and northern parts of the species’ distribution.

## Results

### Thermal performance of *O. tshawytscha* reflects historic stream temperatures

*O. tshawytscha* gametes were sourced from Ship Creek in Anchorage Alaska (AK), Medvejie hatchery in Sitka Alaska (SK), Warm Springs River in Oregon (OR) and the Lower Crab Creek in Washington (WA). Mean summer temperatures during the warmest month of the year at these sites are 8.53°C (AK), 8.46°C (SK), 15.03°C (OR) and 20.56°C (WA). Thermal performance was characterized for each population by determining the maximum metabolic rate (MMR), routine metabolic rate (RMR), absolute aerobic scope (AAS), critical oxygen tension (P_crit_) and acute hypoxic tolerance (LOE hypoxia) after 24h semi-acute exposure to temperatures ranging from 4-24.5°C. All collected metrics of thermal performance varied with both temperature and population but not their interaction, except for LOE hypoxia (Figure 1a,b, Supplementals 1-3). Population-specific patterns were most apparent for AAS, where populations from warmer source environments (OR and WA) displayed AAS curves that were shifted to warmer temperatures compared to AK or SK populations (Figure 1a). The shift in AAS was driven by a reduction in RMR at high temperatures in the OR and WA fish (Figure 1b). Similarly, thermal breadth was several degrees higher in the warm adapted populations (Figure 1b). The reduced routine metabolic demand of the OR and WA populations likely underlies the improved hypoxia tolerance of these populations (S2&3) demonstrating a protective effect of reduced metabolic demand during periods of low oxygen availability. Notably, the thermal optima (T_opt_) for AAS varied from 17.9°C in AK fish to as high as 24.6°C in the WA fish (Figure 1a). The WA population was particularly notable as the predicted T_opt_ of 24.5°C is well above the T_opt_ usually observed in this species (Poletto et al., 2017; Zillig, Lusardi, et al., 2023), which may reflect adaption to the thermally challenging stream in which this population spawns, where summer temperatures commonly exceed 26°C (Small et al., 2011) similar to the summer temperatures seen at the extreme southern end of this species range (Zillig, FitzGerald, et al., 2023).

**Figure 1:**
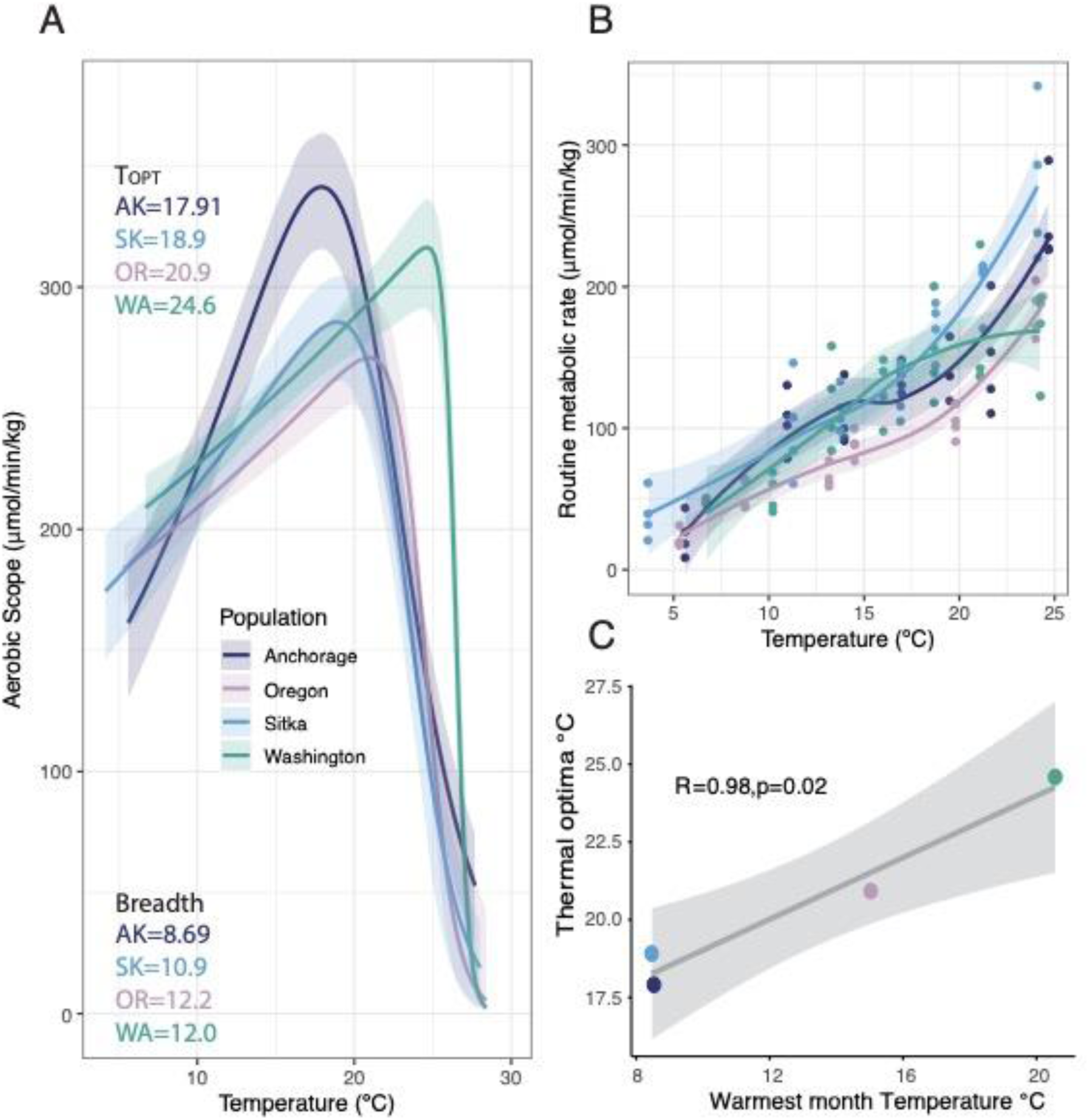
Aerobic performance in Chinook salmon is locally adapted: A) Thermal performance curves generated from absolute aerobic scope for each population with calculated values for thermal optima (T_opt_) and thermal breadth (temperature range across which 80% aerobic scope is maintained). Confidence intervals are derived from residual bootstrapping. B) Smoothed routine metabolic rate data colored by population where dots are individual fish. C) Correlation between calculated population-specific thermal optima and the average temperature of the warmest month of the year from the location where each population was sourced from.

The T_opt_ for the various salmon populations displayed strong correlation with historical average stream temperature during the warmest month of the year (Figure 1c). These results indicate that even when reared under identical temperatures in the lab, population-specific respiratory performance is maintained, providing evidence for genetically based local thermal adaptation. Interestingly, while the T_opt_ for each population is above the mean summer temperature, this safety margin is higher in the cool-adapted populations indicating a potentially greater resilience to climate warming in the Northern parts of this species range.

### Population Physiotype Differentiates Warm Versus Cool-adapted Populations

As a further measure of thermal performance, we were interested in how the observed population differences in aerobic activity would correspond with acute thermal tolerance (CT_max_). CT_max_ varied by population and mirrored the patterns of AAS, with OR and WA fish having improved acute thermal tolerance compared to the AK and SK fish (Figure 2a). Interestingly, the difference in CT_max_ between populations was small (∼1°C) compared to the differences in T_opt_ (∼7°C). One potential explanation for this difference is the “concrete ceilings and plastic floors hypothesis” (Sandblom et al., 2016) which states that absolute thermal limits like CT_max_ are constrained by biochemical processes, while setpoints like RMR are more flexible and thus modulation of these setpoints influences thermal performance. The similar patterns displayed by CT_max_ and AAS could either arise if these traits are mechanistically linked, or if these traits are independent but shaped by the same selective forces.

**Figure 2:**
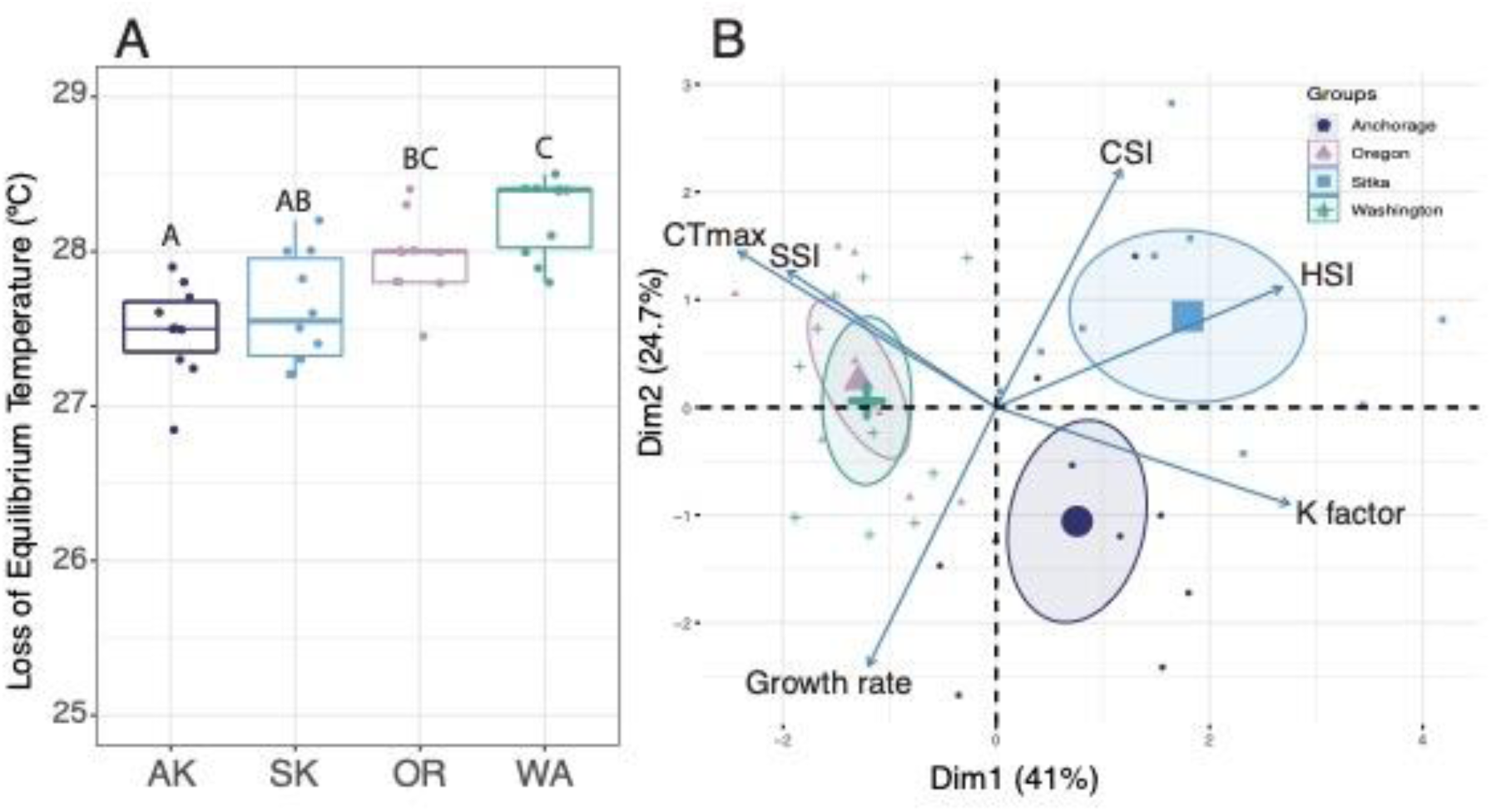
Population-specific physiotypes: A) Average CT_max._ from each population where individuals are represented by data points. Letters above boxplots indicate significant differences between groups with a one-way anova following Tukeys HSD correction (n=10/group). B) Scaled PCA showing phenotypes for fish in CT_max_ trials. Ellipses denote 95% confidence level for a multi-variate t-distribution and samples are color coded by population.

In addition to CT_max_, body shape, growth rate, hepatosomatic index (HIS) and spleenosomatic index (SSI) likewise display populations level differences (Figure 2a, b and S3-7). These phenotypic differences demonstrate a warm-adapted physiotype displayed by the OR and WA fish characterized by larger spleens and a cool-adapted physiotype displayed by the AK and SK fish characterized by a longer body shape and larger livers. The larger relatively spleen size observed in the warm adapted populations is interesting as thermal tolerance in Salmonids has been ascribed to the ability to maintain cardiorespiratory function as temperatures rise (Eliason et al., 2013) and in fish, the spleen serves as an erythrocyte reservoir (Joyce & Axelsson, 2021). Previously, hematocrit was shown to be associated with CT_max_ in *O. tshawytscha* (Muñoz et al., 2018), and thus larger spleens may be an axis of differentiation which facilitates adaptation to warmer environments by improving cardiorespiratory capacity through having an increased erythrocyte reservoir.

### Expression of genes associated with CT_max_ are under evolutionary constraint and enriched for respiratory function

To gain insight into the potential mechanistic basis of acute thermal tolerance in *O. tshawytscha* we performed mRNAseq on ventricle tissue from fish that underwent CT_max_ trials compared to controls held at the rearing temperature of 10°C. Gene expression profiles were dominated by treatment condition with source population accounting for a smaller proportion of variation (Figure 3a). Overall, the gene expression response to CT_max_ was highly similar between populations (Figure 3b) and was enriched for numerous processes related to thermal stress including: protein folding, response to heat, and response to ischemia. Our observed gene expression patterns mirrors previous work which identified reduced oxygen availability in the heart from venous return and subsequent disruption of cardiac activity as a potential physiological basis of thermal limits in *O. tshawytscha* (Clark et al., 2008).

**Figure 3:**
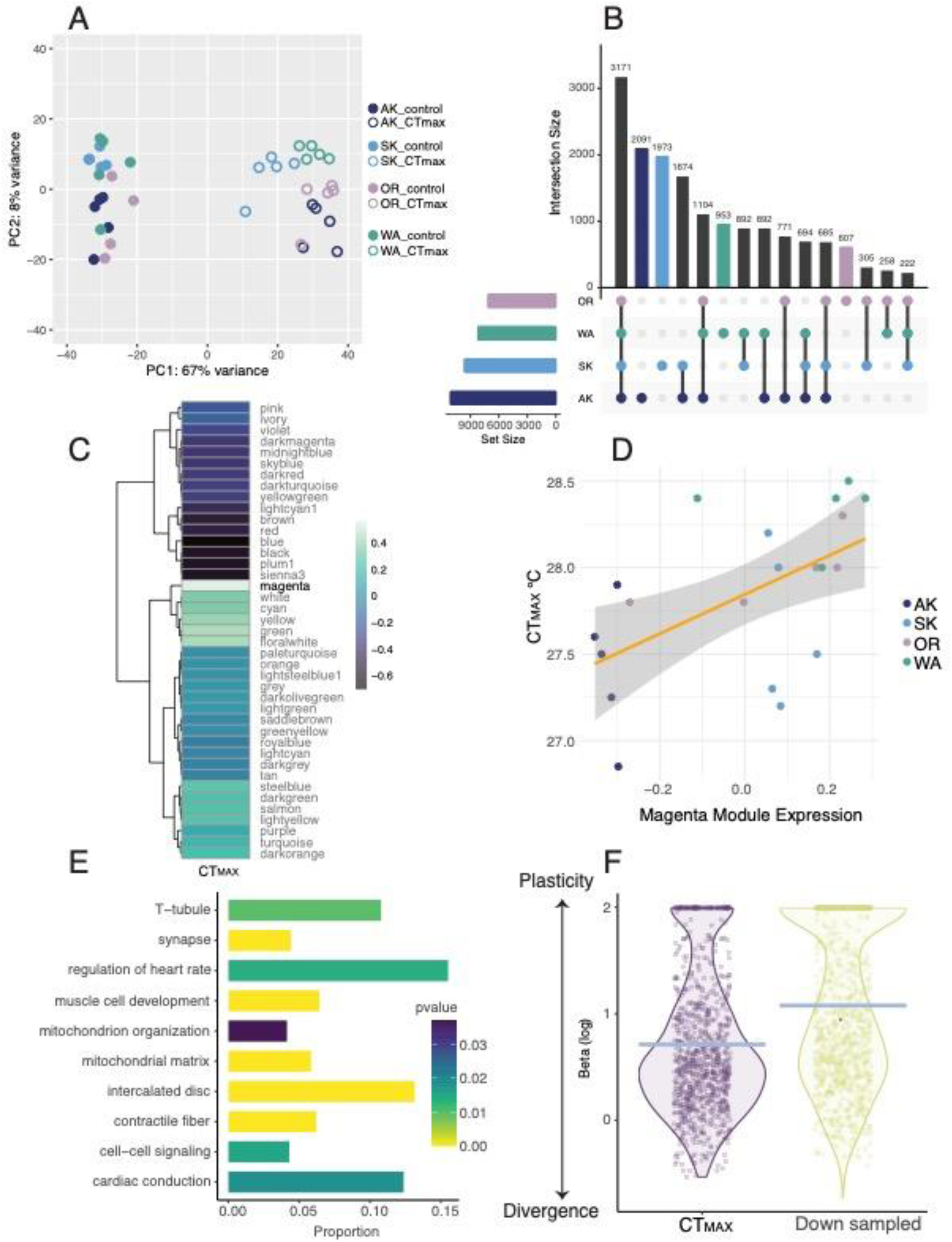
Genetic basis of thermal tolerance. A) PCA of gene expression profiles from all samples, where populations are color coded. Expression profiles of Control versus CT_max_ fish are differentiated with open versus filled circles. B) Upset plot showing the overlap in differentially expressed genes as a consequence of exposure to CT_max_ between all population combination. C) Heatmap showing WGCNA module-trait correlations to CT_max_. D) Correlation between the expression of the WGCNA module associated with CT_max_ (magenta module) and fish CT_max_ E) Barplot showing selected top GO enrichments of the 926 genes in the magenta module. Color denotes FDR-corrected pvalue, while proportion denotes the percentage of genes (transcriptome-wide) for term which are in the magenta module. F) Log-normalized Beta values calculated from the EVE model for the genes in the CT_max_ module compared to a size-matched random down sampling of genes. Blue line indicates mean which is significantly different (pvalue < 2.2e^-16, two-tailed t-test). Lower beta values represent expression divergence of the gene between populations, while higher beta values represent greater expression plasticity of the gene. The beta value is a ratio and thus a unitless metric.

To gain further insights into the genetic basis of population specific thermal limits we performed Weighted Gene Co-expression Network Analysis (WGCNA) (Langfelder & Horvath, 2008) on the transcriptomic data of CT_max_ treated fish. This analysis revealed one module (magenta) whose expression is highly associated with CT_max_ (Figure 3c,d; R2=0.54, p=0.015) that contains 926 genes enriched for terms involved in cardiac contraction and mitochondrial function (Figure 3e). Notably, the individual gene with the highest correlation to CT_max_ in the magenta module was the nuclear encoded mitochondrial respiratory subunit Cytochrome Oxidase 7b (R2 = 0.84, p = 0.001; S9). Overall, these data indicate that variation in CT_max_ is tightly associated with the expression of genes related to ventricular contraction and mitochondrial function. These results support work in other salmonids which demonstrate that reducing cardiac perfusion via coronary ligation reduces CT_max_ (Morgenroth et al., 2021), suggesting that the upper thermal limits of *O. tshawytscha* may be determined by the point at which oxygen availability in the heart, necessary to support cardiac output, becomes limiting.

As our respirometry data suggest that AAS demonstrates patterns consistent with local adaptation, we employed the evolutionary analysis of variance model (EVE) (Gillard et al., 2021; Rohlfs & Nielsen, 2015) to identify genes which display localized expression profiles indicative of selection. We identified 254 genes displaying putatively adaptive expression patterns (FDR < 0.1). To assess the role of thermal tolerance as a driver of selection we compared the 254 genes displaying adaptive expression patterns with the 926 genes in the WGCNA module associated with CT_max_. Overall, we found high overlap between these two independent gene sets as the putatively adaptive genes were nearly three times as likely as a size-matched random down sampling of genes to be in the CT_max_ associated module (p-value = 0.00476). Furthermore, the CT_max_ associated genes on average displayed more divergent expression profiles than these down-sampled genes (p-value < 2.2e^16; Figure 3f). Taken together the results suggest that the expression of genes associated with CT_max_, specifically those involved in mitochondrial energy production and ventricular contraction, are more likely to develop localized expression patterns, indicating that the expression level of these ventricular genes is under strong localized selective pressure.

Overall, our data indicate that the elevated thermal tolerance of the OR and WA fish is not driven by the activation of unique pathways in response to temperature stress, but rather different expression setpoints of metabolic and contractile genes. These gene expression patterns in tandem with our observations of population-level variation in RMR at high temperatures support selection of metabolic setpoints as a mechanism of thermal adaptation consistent with the “concrete ceilings and plastic floors” hypothesis (Sandblom et al., 2016). Overall, we demonstrate multiple lines of evidence which indicate that population-level differences in the expression of mitochondrial genes play an important role in thermal performance of *O. tshawytscha*.

### Thermal responsiveness of ventricular mitochondria differentiates warm versus cool-adapted populations

As our gene expression data highlighted the expression of mitochondrial genes as a potential contributor to localized thermal performance, we sought to identify how the mitochondrial physiology of ventricle tissue varies by population both at a permissive temperature (16°C) and near whole organism CT_max_ (27.5°C). At 16°C neither mitochondrial oxygen consumption rate (OCR) nor proton efflux rate (PER), a proxy for glycolysis, varied by population (p>0.05; S10&11). However, the thermal responsiveness of both measures of mitochondrial physiology varied by population (Figure 4a & b; p < 0.002) and displayed a clear trend where the cool-adapted populations increased both OCR and PER while the warm-adapted populations showed either a slight decrease or no change. This demonstrates that under near-ideal temperatures, ventricular mitochondria function similarly between populations and that population-level differences only arise at high temperatures, in the form of reduced mitochondrial responsiveness to temperature in the warm-adapted populations. Surprisingly, mitochondrial efficiency as assessed via the respiratory control ratio (RCR) increased from 16°C to 27.5°C (p< 0.05) but did not vary by population (S12), indicating that at this high temperature mitochondria remain functional. While the trends in RCR are consistent across populations our observed values for RCR are lower than those observed in other species of fish (Chung & Schulte, 2020; Iftikar & Hickey, 2013), likely owing to incomplete pharmacological inhibition of ATP synthase during our assay and therefore these data should be interpreted in a relative not absolute sense. Taken together, our data suggest that at supra-optimal temperatures, ventricular mitochondria remain functional and require a greater amount of oxygen in cool-adapted but not warm-adapted populations. At the organismal level, this indicates that when exposed to high temperatures the oxygen demand of the ventricle in warm-adapted populations is lower than in cool adapted populations allowing for continued ventricular pumping. Our findings support previous work demonstrating that mitochondria often remain functional at temperatures above organismal CT_max_ (Chung & Schulte, 2020), and suggests that the role of mitochondria in setting thermal limits stems from thermal responsiveness rather than functional collapse of the organelle.

**Figure 4:**
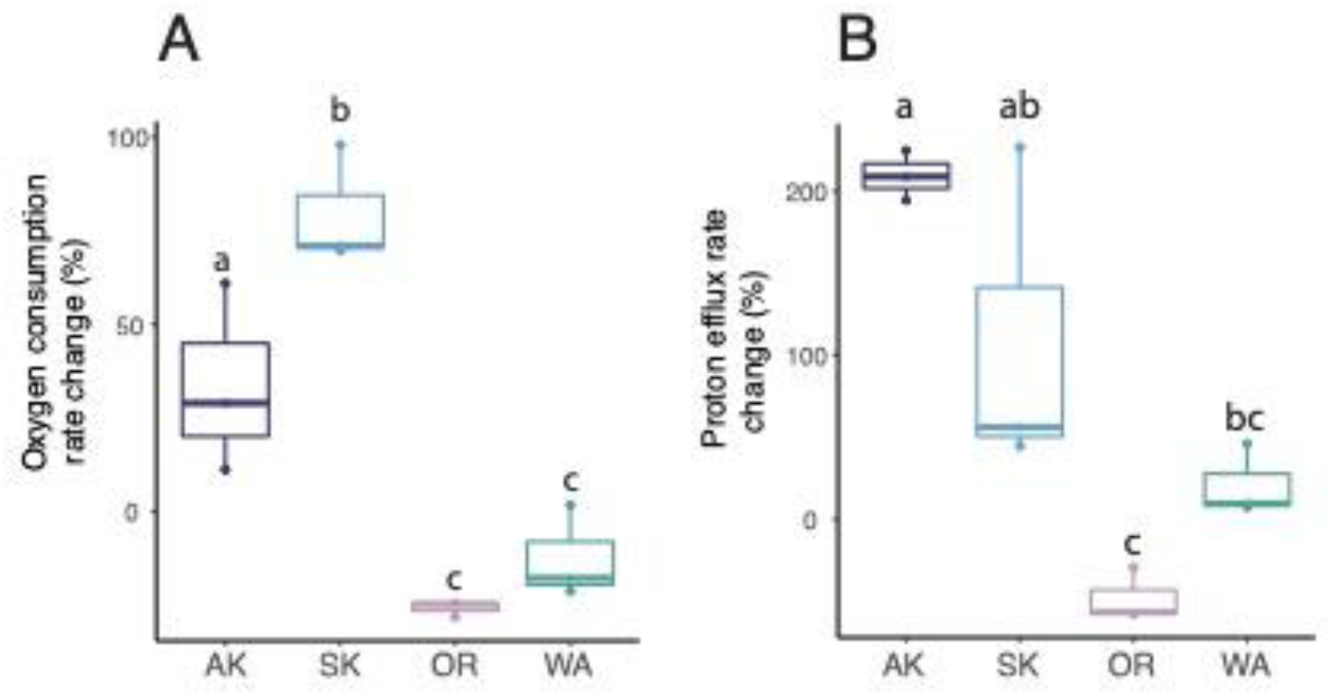
Mitochondrial function varies by population. A) Mitochondrial oxygen consumption rate of change from 16 to 27.5 °C for each population where individuals are represented by dots. Differences between populations following post-hoc testing are denoted by lettering. B) Proton efflux rate (glycolysis) rate of change from 16 to 27.5 °C for each population where individuals are represented by dots. Differences between populations following post-hoc testing are denoted by lettering.

It is worth noting is that 27.5°C is at the CT_max_ of both AK and SK populations but lower than the CT_max_ of either the OR or WA populations. Thus, CT_max_ may represent the temperature at which the energy required to support ventricular activity exceeds the ability of the cell to produce energy via oxidative phosphorylation and glycolysis. From an organismal standpoint an inability to meet the energetic demand of ventricular cardiomyocytes would lead to loss of ventricular contraction and collapse of the cardio-respiratory system akin to a heart attack. In fish, the heart is the last organ to receive oxygenated blood and it relies primarily on cardiac circulation with coronary circulation providing an accessory role (Farrell, 2002). Thus, as the metabolic demands of the systemic tissues rise along with increasing temperatures, having reduced thermal responsiveness in ventricular mitochondria as seen in the OR and WA fish may confer improved thermal tolerance by extending the temperatures at which the heart can continue to circulate blood.

Our data demonstrate that mitochondrial functional traits, whole organism respiratory physiology, as well as gene expression patterns all point to a key axis of thermal adaptation acting at the level of the mitochondria. The locally adapted peaks in AAS are driven by a combination of population-specific metabolic setpoints and lower thermal responsiveness in the warm-adapted populations, which is reflected in population-specific expression of mitochondrial genes in ventricle tissue. Overall, our results indicate that the thermal responsiveness of mitochondria is a key mechanism through which *O. tshawytscha* adapt to thermal environments with effects cascading up from the level of the organelle to the whole organism.

## Discussion

Understanding the genetic, cellular and physiological mechanisms that influence thermal tolerance of populations is critical for assessing vulnerability to rising temperatures. Our results demonstrate that in *O. tshawytscha,* modulating the thermal responsiveness of respiratory capacity is a mechanism through which populations adapt to local stream conditions. The development of locally optimized expression levels of key genes involved in respiration underlies the thermal responsiveness of ventricular mitochondria and thus determines at which temperatures the cardiorespiratory system is able to operate and continue distributing oxygen to the tissues. Based on this mechanistic understanding, our results indicate that the thermal adaptability of *O. tshawytscha* populations is well matched to current temperatures. Additionally, as populations of *O. tshawytscha* are in decline throughout their range, including locations where summer temperatures are well below those which cause physiological impairment the region-wide decline of *O. tshawytscha* and failure to recover is unlikely to be the result of the effect of rising stream temperatures experienced by fish during freshwater rearing.

## Methods and Results

### Fish rearing

Fish used in this study were collected from the William Jack Hernandez sport fish hatchery in Anchorage, Alaska (AK), the Medvejie hatchery near Sitka, Alaska (SK), the Warm Spring National fish hatchery in Warm Springs, Oregon (OR) and the Lower Crab Creek near Royal City, Washington (WA) under permit numbers (AK and SK (Collection SF2021-190, Transport: 8597-06-10-21 and 8596-06-10-21), OR (Transport: 8599-06-07-21), WA (Collection: Lopez 21-255, Transport: 8600-06-15-21). Gametes from AK, SK and WA fish were collected on-site and transported back to Washington State University (WSU) Pullman, WA, USA to be fertilized, while the OR fish were fertilized on-site at the hatchery and were transferred to WSU as eyed-eggs (∼2 weeks post-fertilization). Fertilization dates were as follows: AK-8/9/2021, SK-8/20/21, OR-10/25/21, WA-1/14/22. AK, SK and WA fish originate from fall-run stocks while the OR fish originate from a spring-run stock. All fish were reared under identical conditions, although the exact dates differ by population as these populations all spawn at slightly different times. River temperature data were collected for the Washington and Oregon sites from the NorWeST stream temperature map which uses averages of the mean August stream temperature from 1993-2011. As the NorWeST stream temperature map does not collect data from Alaska we collected data from this same period from USGS gage stations located near the hatcheries at gage stations AK – 15276000 and AK-15087810.

Eggs were incubated in standard vertical hatchery egg rearing trays at 10°C until eggs hatched and had absorbed their yolk sacs, upon which fish were transferred to an Aquaneering zRack in 9.5L tanks under a 12:12 light cycle. Families were reared together and included five AK families, four SK families, three OR families, and three WA families where each family was the result of a different cross, so that no parents were shared between families. Fish were fed 1-2 times a day to satiation with BiVita salmon feed (BioOregon) and uneaten food was removed after feeding. When fish were ∼5cm they were elastomer tagged (Northwest Marine Technology) by family and moved to the Thorgaard Center for Salmonid Physiology and Genomics Carver fish hatchery at WSU. Each population was split into three 55L tanks and an equal number of fish from each family was added to achieve a uniform rearing density of 30 fish per tank. The WSU Carver Hatchery uses de-chlorinated municipal water from Pullman, Washington. Chillers were used to maintain the water temperature at 12°C ± 2°C depending on ambient temperatures. Fish were fed 1-2 times once a day to satiation, (BioVita) and were subject to local astronomical light cycle at location (46.7298° N, 117.1817° W). As the fish in each population grew at different rates and we decided to match populations by fork length for phenotype testing, which meant that they differed in age at the time of respirometry testing varying from eight to twelve months post fertilization. Prior to respirometry or CT_max_ testing, fish were moved back into 9.5L tanks in the lab at a density of 4 fish per tank in order to reacclimate fish for two weeks to a 12:12 light cycle, to minimize the influence of the diel cycle on physiology. All experiments were approved by the Washington State University institutional animal care and use committee under protocol #6607.

### Respirometry

Intermittent flow respirometry was conducted using equipment from Loligo systems (Viborg, Denmark). Respirometry trials were conducted in 360mL plexiglass chambers (Loligo system product number: CH24000) where the total respirometry ratio averaged 42:1 water/fish volume. Oxygen consumption data were collected using a 10m flow-through oxygen cell (loligo system product number: OX11220) and transmitted to a witrox four oxygen meter (loligo system product number: OX11875) through a fiber optic cable (loligo system product number: OX11150). Oxygen readings were collected in one-second intervals and processed using the Loligo AutoRespV3 software (Viborg, Denmark). Oxygen probes were calibrated prior to each run in 100% air-saturated water and sodium sulfite. All trials were conducted in an environmental chamber to maintain temperature and the water bath was continuously aerated to maintain 100% air saturation.

For respirometry trials, fish were exposed to the test temperature by placing fish in the environmental chamber 24h prior to the start of the respirometry trial in 18L aerated tanks at 10°C which were placed in environmental chambers and allowed to reach chamber temperature. Fish were fasted during the acclimation process to minimize the effects of digestion on oxygen usage during the trial. Trials were conducted at seven temperatures per population (7°C, 10 °C, 13°C, 16°, 19°C, 22°C, 25°C). Fish were deprived of food for the 24 hours prior to respirometry trial. Because trails were run over the course of several months trials were conducted in the following order: 10°C, 16°C, 22°C, 13°C, 19°C, 25°C, 7°C to ensure that both high and low temperature trials were conduced both early and late in the series. We additionally attempted to run 28°C trials, however the fish were unable to successfully acclimate at this temperature. Actual temperatures varied somewhat from the goal temperatures during the trial due to limitations in the environmental chamber but varied by approximately 1°C within a trial. The coldest temperature measured varied more but was never greater than 1.5°C as maintaining the lowest acclimation temperature necessitated the use of a recirculation line run through an ice bath which made it more difficult to precisely to control the temperature setpoint. Temperature was recorded continuously with the loligo temperature probe and for analysis we used the average temperature over the 24h+ respirometry trial.

In order to measure aerobic scope, we collected both maximum metabolic rate (MMR) and routine metabolic rate (RMR). For MMR measurements fish underwent a Ucrit swimming protocol in a 51L swim tunnel (Loligo system product number: SW20020), where fish were allowed to acclimate to the swim tunnel for 15 minutes prior to undergoing the swim challenge. The trial began by swimming the fish together at approximately 1.5 body lengths/second (owing to natural variation in fork length between individuals) for 30 minutes. Following this initial velocity, the speed was increased by 0.5 body lengths/second every ten minutes until the fish was pushed against the back grating and could not be motivated to continue swimming via prodding. After fish were exercised to exhaustion, they were immediately placed in respirometry chambers and trials began. Our intermittent flow respirometry protocol involved six-minute measurement periods, one-minute flush periods and one-minute wait periods. On average less than five minutes elapsed between the end of the swimming trial and the start of respiration measurements. The intermittent flow respirometry protocol was kept consistent throughout the course of the experiment and was chosen based on preliminary work to ensure that the measurement windows were long enough to allow the fish to deplete a sufficient amount of oxygen to collect a reliable signal even at the colder temperatures but were short enough to avoid oxygen saturation dipping below 75% at the high temperatures. Trials were conducted for a minimum of 24 hours to allow fish to reach RMR, and after placing fish in the chambers the lights were turned out and door locked to ensure minimal disturbance to the fish. While it has been suggested that periods longer than 24 hours are required to allow fish to achieve their “true” minimum or standard metabolic rate, we use the term routine metabolic rate (RMR) to acknowledge that, given our experimental protocol, fish may have not had sufficient time to achieve their true minimum metabolic rate and that the chambers were large enough to allow small amounts of spontaneous activity (Clark et al., 2013). However, it generally took individuals around 8 hours until their first measurement window reached their RMR after which metabolic rate oscillated around this value. Two background measurements were taken before the fish was placed in the respiration chamber and two background measurements were taken after the conclusion of the experiment in order to account for background respiration.

In addition to collecting respiration data, we also collected information on hypoxia tolerance by halting flush cycles and allowing the fish to deplete the oxygen in the chamber. Hypoxia trials were run for trials above 10°C (for colder trials oxygen usage was low enough that allowing the fish to deplete oxygen was not feasible). Two measurements of hypoxia tolerance were collected, loss of equilibrium (LOE) and critical oxygen pressure (Pcrit). Hypoxia LOE was recorded as the oxygen pressure where fish lost equilibrium and were unable to right themselves for a minimum of ten seconds. Pcrit was collected during data processing described in more detail below. At the end of the trial, fish were recovered in fully saturated water and fork length as well as weight were determined. On average, fish were 92.1mm ± 15.5mm. Fish on average weighed 9.83g ± 5.14g. Four fish from each population were tested at each temperature, for an N of 112 individuals, however five individuals had to be removed from analysis due to data irregularities such as equipment malfunction for a total of 107 fish analyzed. No fish were used in more than one trial to avoid potential impacts of acclimation. Due to differences in growth rate and hatching times fish trials were conducted over the course of several months. Trials for AK fish occurred during May 2022, trials for SK fish occurred in June 2022, trials OR fish occurred in November 2022 and trials for WA fish occurred in February 2023. Trials for each population were conducted over the course of sixteen consecutive days.

### Respirometry Analysis

Respirometry data were analyzed using the R package RespR (Harianto et al., 2019) following the intermittent flow vignette. For this analysis the raw oxygen pressure in kPa is divided by flush, wait and measurement windows. Oxygen consumption is averaged over the length of the window and windows are only retained if the regression coefficient between oxygen levels and time is greater than 0.95. To account for the role of background respiration due to bacterial growth or other processes we used the linear background correction method. This method assumes a linear rate of increase in background respiration based on the pre- and post-trial background measurements and applies a customized correction for each window based on time. MMR was determined as the measurement window with the highest overall rate of oxygen usage (MO_2_), which had a reliable oxygen trace upon visual inspection. The MMR measurement was generally within the first 20 windows. It is not uncommon for the MMR to be outside of the first window as during exhaustive exercise fish often accumulate an oxygen debt due to glycolysis which needs to paid off over time (Clark et al., 2012). Individual RMR was calculated using the fishMO_2_ R package (Chabot et al., 2016) where RMR was called as the average of the lowest 10% of values. AAS is calculated as MMR-RMR. All values of MMR and RMR used to calculate AAS are within the range of values previously observed in juvenile salmonids (Poletto et al., 2017; Zillig et al., 2023). Pcrit was calculated via the (oxy_crit) function in RespR using the segmental method applied to the oxygen data during the hypoxia trial using a width of 0.2. This function works by calling Pcrit as the oxygen pressure where metabolic rate undergoes a clear change in slope (to represent the onset of the hypoxic response). For both measures of hypoxia tolerance there were numerous missing values (30 for Pcrit and 42 for LOE) due to either an inability to identify a reliable breakpoint in the oxygen depletion data or abnormal fish behavior prior to loss of equilibrium.

Each response variable (MMR, RMR, AAS, Pcrit, LOE) was tested separately using a two-way ANOVA with formula *respirometry parameters* (*MME*, *etc*.)∼*temperautre* ∗ *population*. MMR varied both by temperature (p= 5.53e-13) and population (p=2.47e^-5) but not their interaction (p =0.205). (S1). RMR varied by temperature (p< 2.2e^-16) and population (p=9.01 e^-5) but not their interaction (p=0.299) (Figure 1). AAS varied by temperature (p=5.61e6-05) and population (p=0.0178) but not their interaction (p=0.276) (Figure 1). Pcrit varied by temperature (p=4.42e-14) and population (2.79e^-08) but not their interaction (p=0.255) (S2). LOE varied by temperature (p=8.28e^-10), population (p=1.01 e^-4) and their interaction (p=0.014) (S3).

Thermal performance curves (TPCs) were generated using the R package rTPC (Padfield et al., 2021). Initially AAS curves were run through the model selection script in rTPC which fits 25 TPC model formulations to the data with the best model being selected via AIC. As the Washington population experienced no decline in aerobic scope at the highest recorded temperature 24.6°C we elected to constrain the upper bounds of the model by using the observed CT_max_ data from each population to set the point at which each population AAS crosses the x-axis. When analyzing the aggregated AAS data from the four populations the pawar_2018 model (Kontopoulos et al., 2018) was selected as the best model based on AIC. The rTPC model selection script provides AIC, AICc and BIC values, however the model scoring order was not substantially different regardless of model scoring method. The AAS data from each population were fit separately to the pawar_2018 model to generate predicted AAS values with confidence intervals as well as respirometry parameters (Topt, Rmax, Tcrit, etc.) by conducting residual resampling bootstrapping over 500 iterations.

### Acute Thermal Limit (CT_max_) trials and sampling

Two weeks prior to CT_max_ testing fish were moved to 9.5L tanks in an Aquaneering zRack on a 12:12 light cycle in order to minimize the influence of diel cycle on CT_max_. Trials were conducted in a custom built CT_max_ apparatus where fish were suspended in mesh bags in a clear sided 75L container that was aerated and connected to a header tank containing heaters. The header tank was used to heat water at a constant rate of 0.1°C/minute and a 30L/min water pump was used to ensure sufficient water mixing. Fish were deprived of food for 24 hours prior to the start of the trial to reduce the metabolic demands of digestion. CTmax trials lasted ∼180 minutes as water was heated from a starting point of 10°C. CT_max_ values were recorded as the point at which fish lost equilibrium and were unable to right themselves for 10 seconds. Temperature was recorded at three points in the container and the thermometer closest to the fish was used to record CT_max_ values at a resolution of 0.1°C. CT_max_ values for AK fish averaged 27.49°C ± 0.303°C. SK fish averaged 27.62°C ± 0.363°C. OR fish averaged 27.96°C ± 0.269°C. WA fish averaged 28.23°C ± 0.254°C. CT_max_ values were compared with a one-way ANOVA *CTmax*∼*population* which was significant (p = 1.03 e^-5). Differences between populations were assessed via a TukeyHSD which revealed significant differences between (OR-AK, WA-AK and WA-SK at p< 0.05 and between OR-SK at p < 0.1).

Once a fish lost equilibrium it was immediately transferred to a bath containing a lethal dose of MS-222 and were killed with a blow to the head. The fish were weighed, measured for fork length, photographed and checked for elastomer tags. Unfortunately, elastomer tags had fallen out of most fish meaning that we are unable to incorporate the role of family in our models. The liver, heart and spleen were removed from each fish and washed quickly in a bath of phosphate buffered saline (VWR product number: P32080-100T). Tissues were blotted dry, weighted and then flash-frozen in liquid nitrogen. The ventricle was isolated with foreceps prior to flash freezing. For control fish to be used in RNAseq we used an identical setup, without heaters and instead added ice in order to maintain water temp at 10°C. These fish underwent an identical dissection protocol as the CT_max_ fish.

All CT_max_ trials began promptly at 10 a.m. and two days of trials were run for each population as the size of the CT_max_ apparatus was limited to five fish. For a total of N=10 CT_max_ fish/population and N=5 control fish/population were sampled, with the exception of Oregon which had only 4 control fish due to limitations in fish number. Fork length averaged 108.75mm ± 13.6mm. Fish weight averaged 14.8g ± 5.11g.

Cardio-somatic index (CSI), spleno-somatic index (SSI) and hepato-somatic index (HSI) were calculated by dividing organ weight by whole fish weight. Fulton’s K factor was calculated following (Froese, 2006) and growth rate was calculated based on FL divided the number of weeks which had elapsed between hatching and CT_max_ testing. Phenotypic differences between populations were assessed via separate one-way ANOVA testing *phenotype*∼*population* which revealed significant associations with SSI (p= 0.008) (S7), growth rate (p=1,21e^-4), HSI (p=8.24e6-7) (S5), K-factor (p=0.00762) (S4) but not CSI (p=0.065) (S6). Multivariate physiotypes were determined via a scaled PCA (Figure 2).

### RNA sequencing and Processing

RNAseq was performed on ventricle tissue for five CT_max_ and five control fish per population. Bulk RNA was extracted with the Zymo Quick-RNA Miniprep Plus kit (product number: R1057) according the manufactures instructions with the addition of a two minutes of glass bead beating in -20°C cooled blocks. Library preparation and sequencing was performed by Novogene. Sample integrity was assessed via RIN values and all samples with RIN > 9 were used for library prep. RNA libraries were prepared via poly-A tailing and sequenced with 150bp, paired-end sequencing on a Novaseq 6000 resulting in an average of 51.3 million reads per sample (std 7.3 million reads) with 96.6% of reads >Q20 and 91.5% > Q30.

Raw reads were cleaned and trimmed using TrimGalore V0.6.6 (https://github.com/FelixKrueger/TrimGalore). Read counts were enumerated with Salmon V1.8.0 (Patro et al., 2017), the salmon index was generated from the *O. tshawytscha* transcriptome (Otsh_v2.0) and decoys were constructed by concatenating the *O. tshawytscha* genome (Otsh_v2.0) (Christensen et al., 2018) to the end of the transcriptome and using the genomic contigs names as decoys. We used a K-mer size of 31 which our previous work has shown to be sufficient for differentiating between transcripts with low sequence divergence such as ohnologs or paralogs which are common in Salmonid genomes (Dimos & Phelps, 2022). Read mapping rates averaged 73.2% resulting in an average coverage of 965 mapped reads/gene. Transcript abundances were summarized to gene level with Tximport (Soneson et al., 2015) to create a count matrix. Genes were annotated using the online portal of eggNOG-mapper v2 (Cantalapiedra et al., 2021) with taxa autodetection.

### Differential Expression testing

PCA revealed no clear outliers, and all samples were retained for analysis. Genes were removed if the row means were below 10 counts. Differential expression testing was performed separately for each population with DESeq2 (Love et al., 2014), testing for the effect of treatment and genes were considered to be differentially expressed at an FDR<0.05 where upregulated genes are those whose expression increases when exposed to acute thermal limits (CT_max_) and down-regulated genes are those whose expression decreases when exposed to acute thermal limits. AK fish had 6275 up-regulated and 6186 down-regulated genes. SK fish had 5085 up-regulated and 5033 down-regulated genes. OR fish had 4744 up-regulated and 3933 down-regulated genes. WA fish had 4874 up-regulated and 4510 down-regulated genes. Of these 1994 genes were upregulated in all populations (S9). Gene Ontology (GO) enrichments were conducted on these commonly upregulated genes using the R script GO_MWU (Wright et al., 2017) with the binary classification using the *O. tshawytscha* genome as the enrichment background. To accomplish this each gene was assigned one if it was contained in the category on which enrichment were being performed (ex. the 3171 commonly differentially expressed genes), or a zero if the gene was not in this list. This method of GO enrichment tests for enrichment of GO terms within the selected set of genes against the genomic background. Identical parameters were used for cellular compartment (CC) and biological process (BP) with smallest = 25, largest = 0.1 and cluster cutheight = 0.25. This method resulted in 31 enriched CC terms and 371 enriched BP terms (FDR<0.05).

### Gene Expression Phenotype Correlations

To identify genes with expression correlated to CT_max_ we employed Weighted Gene Co-expression Network Analysis (WGCNA) (Langfelder & Horvath, 2008). This method uses an all by all correlation matrix between genes to identify groups of genes with corelated expression patterns and groups these genes into modules. The summarized expression of the module as assessed via an eigenvalue that is regressed against a trait of interested (in this case CT_max_) to identify groups of genes whose expression profile is associated with the trait of interest. Raw counts from the CT_max_ fish were vst normalized in DESeq2 by population (Love et al., 2014) and served as the input to WGCNA. Unsigned gene networks were constructed with a power of 9 (based on soft thresholding), a minimum module size of 30, and a merge cutheight of 0.25. This resulted in the construction of 48 modules of which only 1 (magenta module) was significantly positively associated with CT_max_; this module contained 926 genes (R2= 0.56, p= 0.011). GO enrichments were run on the genes in this module with GO_MWU with the modification of using the module membership score (kme) for the genes in the magenta module and performing the unsigned WGCNA module enrichment test within GO_MWU (Wright et al., 2017). This analysis revealed 36 enriched CC terms and 71 enriched BP terms (FDR < 0.05).

### EVE Model

To identify sets of genes which have developed divergent expression patterns between populations we used the updated evolutionary analysis of variance model (EVE model) (Gillard et al., 2021; Rohlfs & Nielsen, 2015). The EVE model examines gene expression as a continuous trait which evolves along a phylogeny by identifying a trait-optima using an Ornstien-Uhlenbeck process. The observed variance from the calculated optima is expressed as a ratio referred to as the beta statistic. Larger beta values denote greater variation within than between populations and represents genes displaying plastic expression patterns, while smaller values of beta denote greater variance between population than within populations and represent genes with divergent expression patterns. Significance is tested for using a chi-square test with one degree of freedom. The input for this analysis is vst normalized counts from all samples and a phylogenetic tree with one tip per population. To create the necessary phylogenetic tree we mapped filtered RNAseq reads from one representative sample per population to the *O. tshawytscha* transcriptome using BWA-MEM2 v2.2.1 (Vasimuddin et al., 2019). We elected to use a representative sample per population as the EVE model only allows one tip per population and each population is represented by a small number of families <= 5. Samtools v1.9 (Li et al., 2009) was then used to filter duplicates, sort and index alignments. Genotype likelihoods were then called using ANGSD v0.921 using the samtools model with a P-value of 1E-6 which identified 4186 high-confidence SNPS. The produced variant call file file was used to generate a UPGMA phylogenetic tree with the package VCF2PopTree (Subramanian et al., 2019). Once the population tree was generated a beta-shared test was conducted in EVE using vst normalized counts from both CT_max_ and control samples. We ensured that model residuals were normally distributed and that the results followed the expected distribution.

Following false discovery rate correction at an FDR of < 0.1 we identified 254 genes displaying divergent expression patterns, of which 28 were contained within the magenta WGCNA model. To test for enrichment against the genomic background we randomly sampled 926 genes and compared the number of genes with divergent expression patterns between the magenta and the random gene set with a two-sided fischer’s exact test which revealed an odds ratio of 2.79 and a p-value of 0.0047, indicating that the significant EVE genes with low beta values were more likely to contained in the CT_max_ associated gene set. To assess the magnitude of the differences we compared the log-normalized beta values between the two gene sets with a two-sided t-test which revealed a significant shift towards lower beta values in the CT_max_ associated genes compared to the randomly selected gene set (pvalue < 2.2e^-16) (Figure 3).

### Mitochondrial Function

An Agilent Seahorse XFp Mito Stress Test kit (Agilent product number: 103016-400) was used to assess mitochondrial function of cell slurries derived from ventricle tissue. In June 2023, Fish were euthanized via a lethal dosage of MS-222 followed by cervical concussion. Hearts were excised and placed in a bath of room temperatures sterile PBS (VWR product number: P32080-100T) where the ventricle was isolated with forceps and efforts were made to remove as much blood as possible from the ventricle. Following isolation, ventricles were placed in a PBS solution containing 10 μl of 20ug/μl of type1 collagenase (VWR catalog number IC195109.1). Ventricle tissue was incubated in collagenase for 15 min at RT on a tube spinner, and a motorized pestle was used to homogenize the tissue every five minutes. Following incubation, the slurries were vigorously mixed with a pipet to minimize tissue clumping. Cells were then centrifuged for 2 minutes at 1000xg and the supernatant was removed. Cells were then washed twice with 10% FBS (VWR product number: 97068-085) in PBS to inactivate the collagenase. After washing, cells were resuspended in Agilent XF-DMEM respiration media (Agilent product number: 103680-100) with 0.01M glucose, 0.1mM pyruvate, and 0.2M glutamine. Protein concentrations of the cell slurries were quantified with a Qubit fluorometer (Fisher product number: Q33211) and wells were seeded with 7.5ug of protein per well, reaching 90-95% confluence. Respiration media was added to wells to achieve 180ul per well. This isolation protocol served to isolate actively respiring ventricle muscle fibers as well as erythrocytes as assessed via TMRE (Fisher product number: T669) positive cells.

The Mito Stress Test kit assesses oxygen consumption rate (OCR) and glycolysis rate in cells under four conditions: resting, ATP synthase inhibition (leak respiration), electron transport chain uncoupling (spare respiratory capacity), and non-mitochondrial respiration (background) using the pharmacological inhibitors oligomycin, FCCP and Rotenone/actimycin, respectively. A pilot test was used to determine concentrations of 3.0M oligomycin, 1.0M FCCP and 1.0M Rotenone. Cells were incubated at two test temperatures 16°C and 27.5°C for thirty minutes prior to the start of the assay. 16°C was chosen to represent near “optimal” conditions for *O. tshawytscha* while 27.5°C was chosen based on the CT_max_ of the least thermally tolerant population (AK). While 27.5°C was a lower temperature than the CT_max_ of the OR and WA populations we were interested in assessing mitochondrial function at a common high temperature without exceeding the thermal limit of the fish these cells were derived from. The Seahorse XFp recorded three measurements per condition, and the values were averaged across the three measurements. Assays were conducted with technical replicates (derived from the same fish) and values were averaged over the two replicates. In the case of extreme values, the oxygen depletion curves were assessed, and if an error occurred that value was removed from the data set. All measurements were conducted over the course of seven days in June 2023. Despite the results from the pilot study, we observed limited efficacy of the injected drugs to cause the desired phenotypic effect, likely due to inefficient drug penetrance or concentrations which were too low to completely inhibit their target. While pharmacological efficiency was below optimal, the expected trends in the data were consistent with the anticipated effect of each drug. Oligomycin reduced OCR, FCCP increased OCR and Rotenone/AA reduced OCR almost entirely. Respiratory control ratio (RCR) which assesses the efficiency of mitochondria was calculated as the basal OCR divided by OCR values following the addition of oligomycin.

All values were normalized to total protein levels and differences between groups were assessed via a two-way ANOVA testing *Mitochondrial parameter* (*OCR*, *PER*, *RCR*)∼*temperature* ∗ *population*. OCR varied by temperature (p=0.0016) and the interaction between temperature and population (p=0.016) but not population (p=0.47) (S11). Glycolysis rate varied by population (p=0.0014), temperature (p=0.036) and their interaction (p=0.0039)(S12). RCR varied by temperature (p=0.012) but not population (p=0.27) or their interaction (p=0.66)(S13). Thermal responsiveness was calculated for each parameter as the difference between the 27.5°C samples compared to the population means observed at 16°C. Thermal responsiveness between populations was compared with a one-way anova testing *Mitochondrial parameter* (*OCR*, *PER*, *RCR*)∼*population* with differences between populations assessed via a TukeysHSD test. Thermal responsiveness of OCR varied by population (p=0.00174) with differences observed between (SK-AK, OR-AK, WA-AK, OR-SK, WA-SK). Thermal responsiveness of PER varied by population (p=0.00187) with differences observed between (OR-AK, WA-AK, OR-SK). Thermal responsiveness of RCR did not vary by population p=0.19.

## Supplementary Figures

**Supplemental Figure 1:**
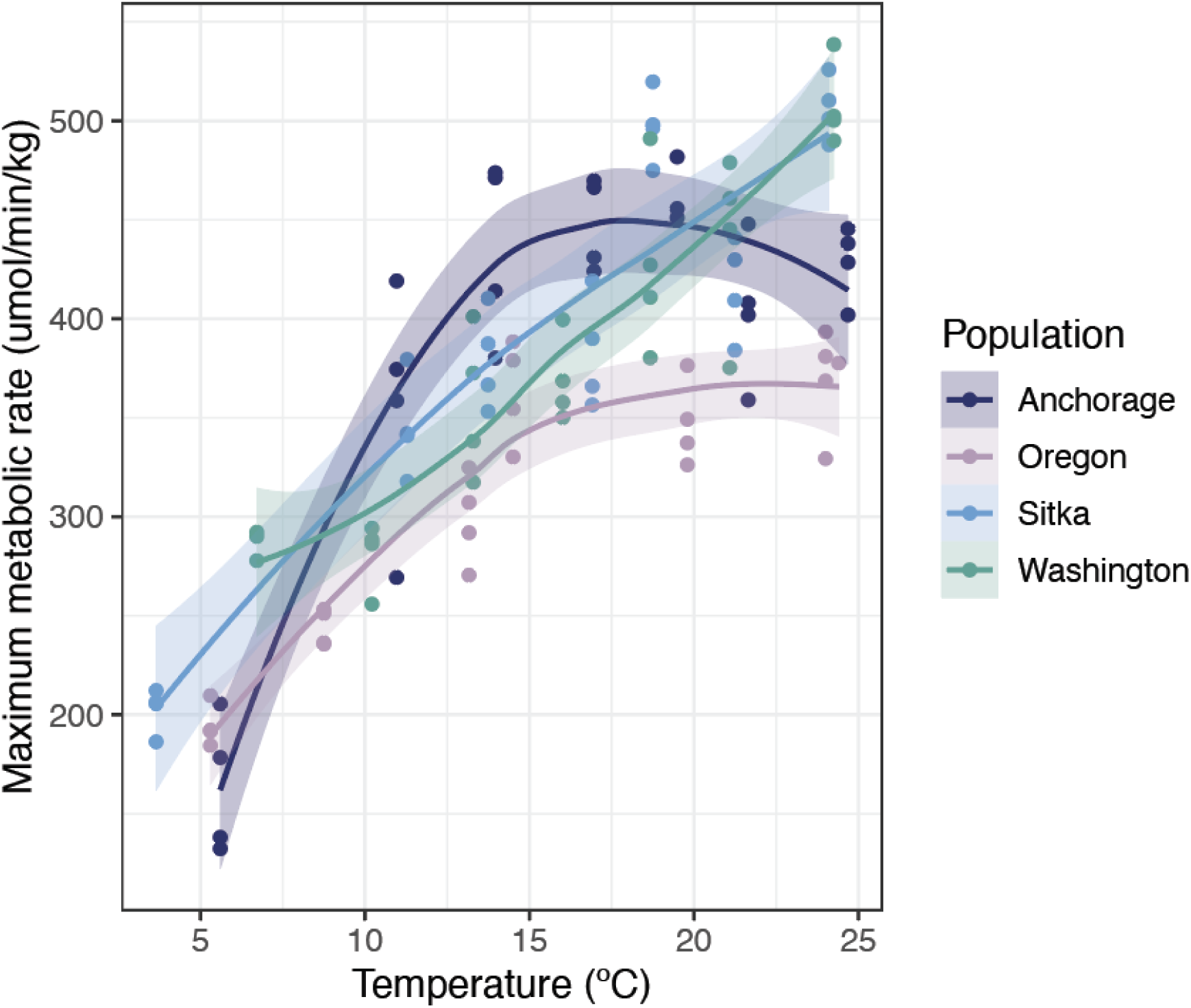
Maximum metabolic rate. Populations are color-coded and individual fish are represented as points.

**Supplemental Figure 2:**
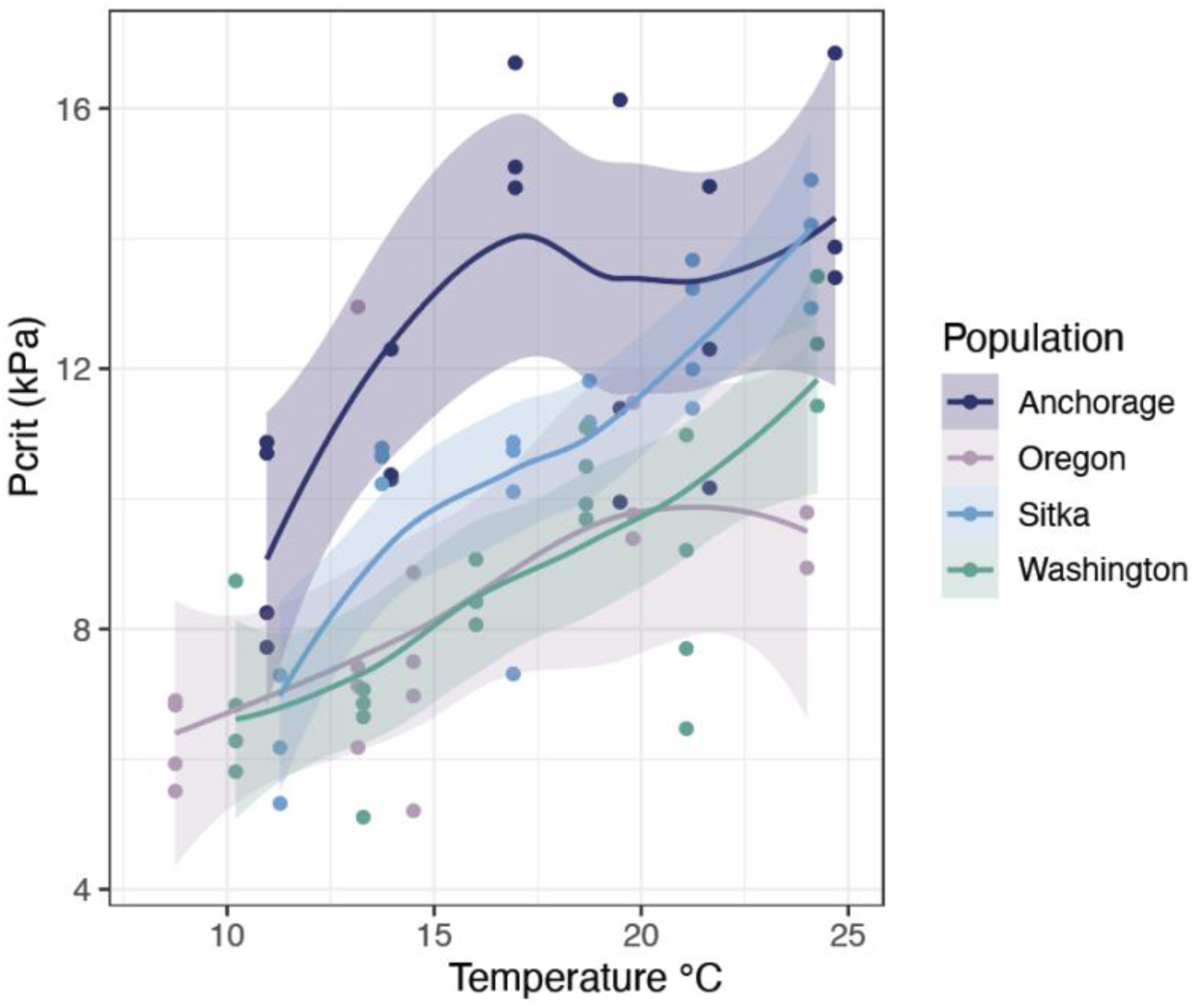
Critical oxygen tension. Populations are color-coded and individual fish are represented as points.

**Supplemental Figure 3:**
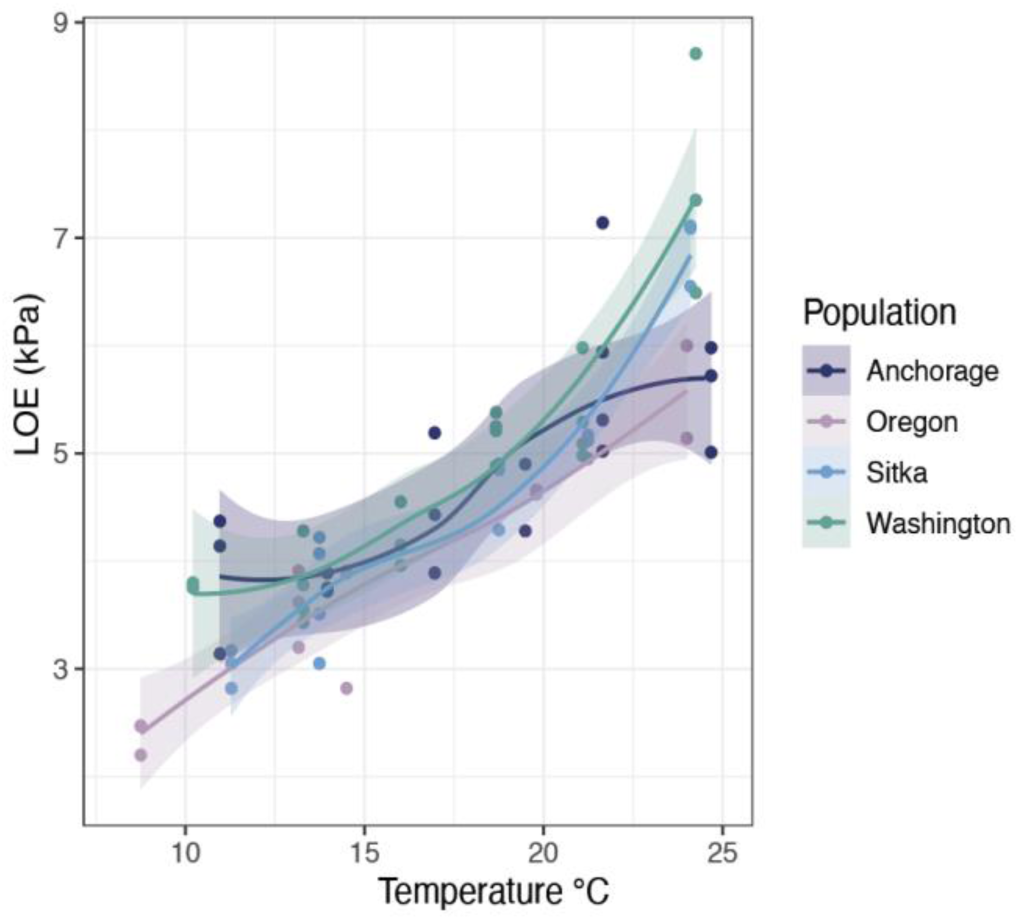
Hypoxia loss of equilibrium. Populations are color-coded and individual fish are represented as points.

**Supplemental Figure 4:**
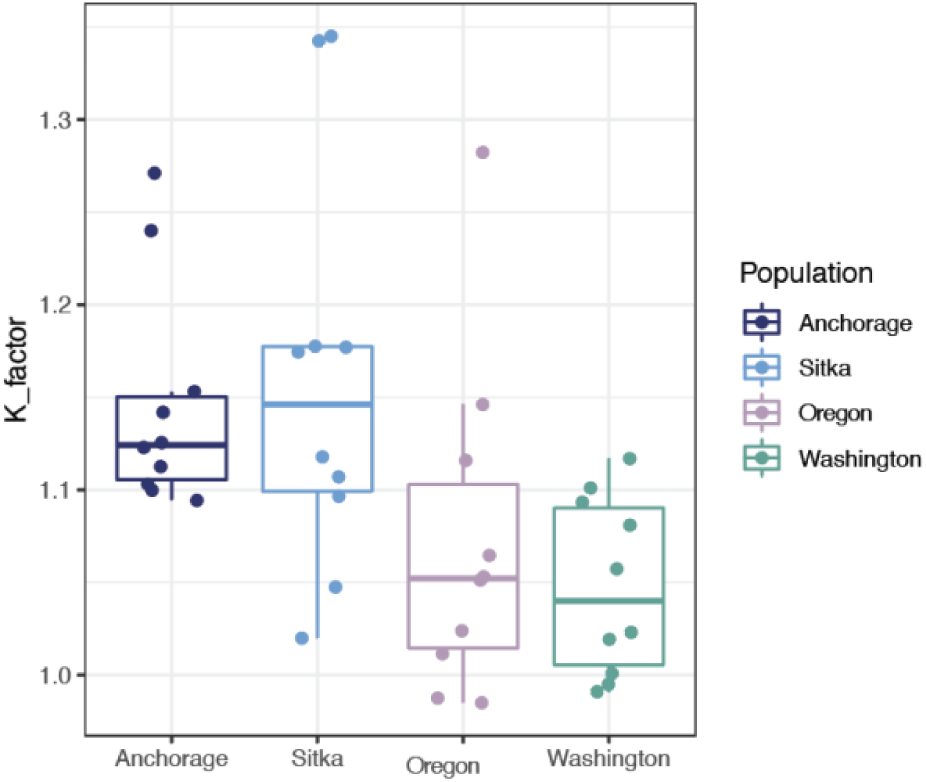
Body conditioning factor represented as K-values for each fish which underwent CT_max_ trials.

**Supplemental Figure 5:**
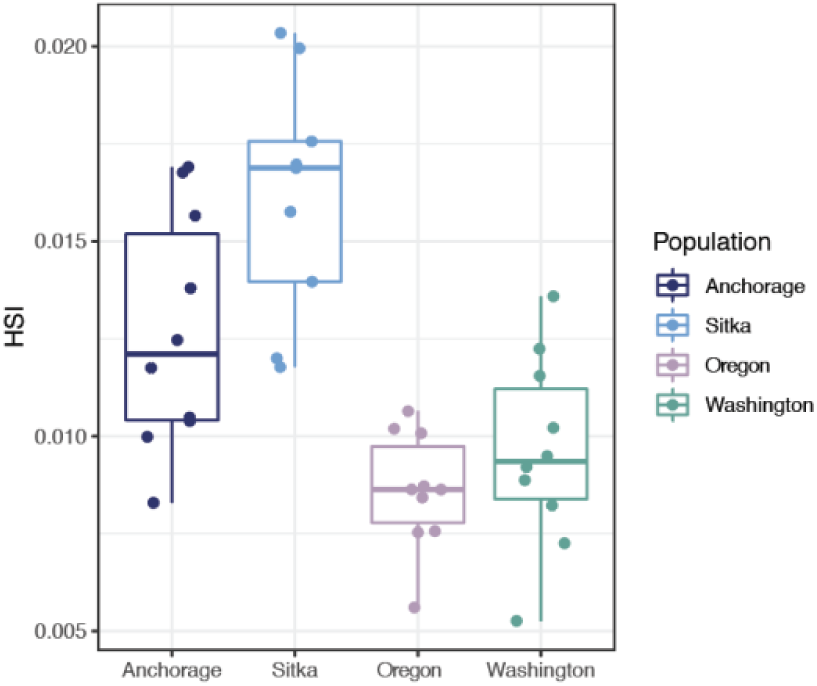
Hepatosomatic index (HSI) for each fish which underwent CT_max_ trials.

**Supplemental Figure 6:**
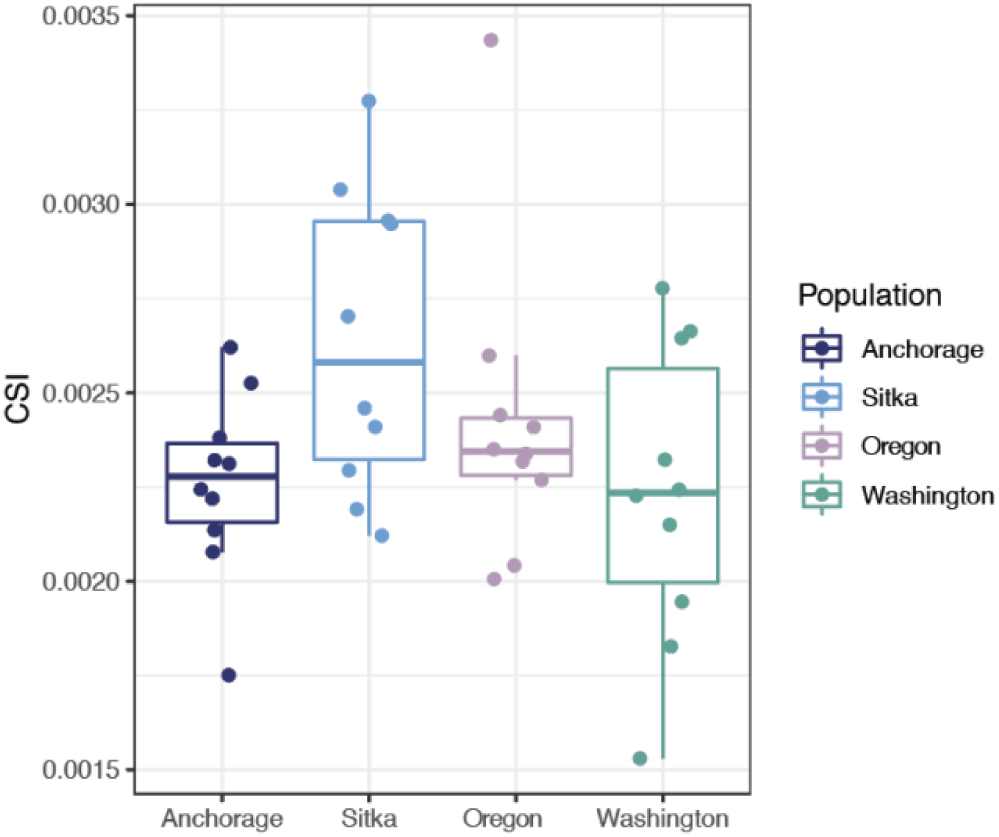
Cardiosomatic index (CSI) for each fish which underwent CT_max_ trials.

**Supplemental Figure 7:**
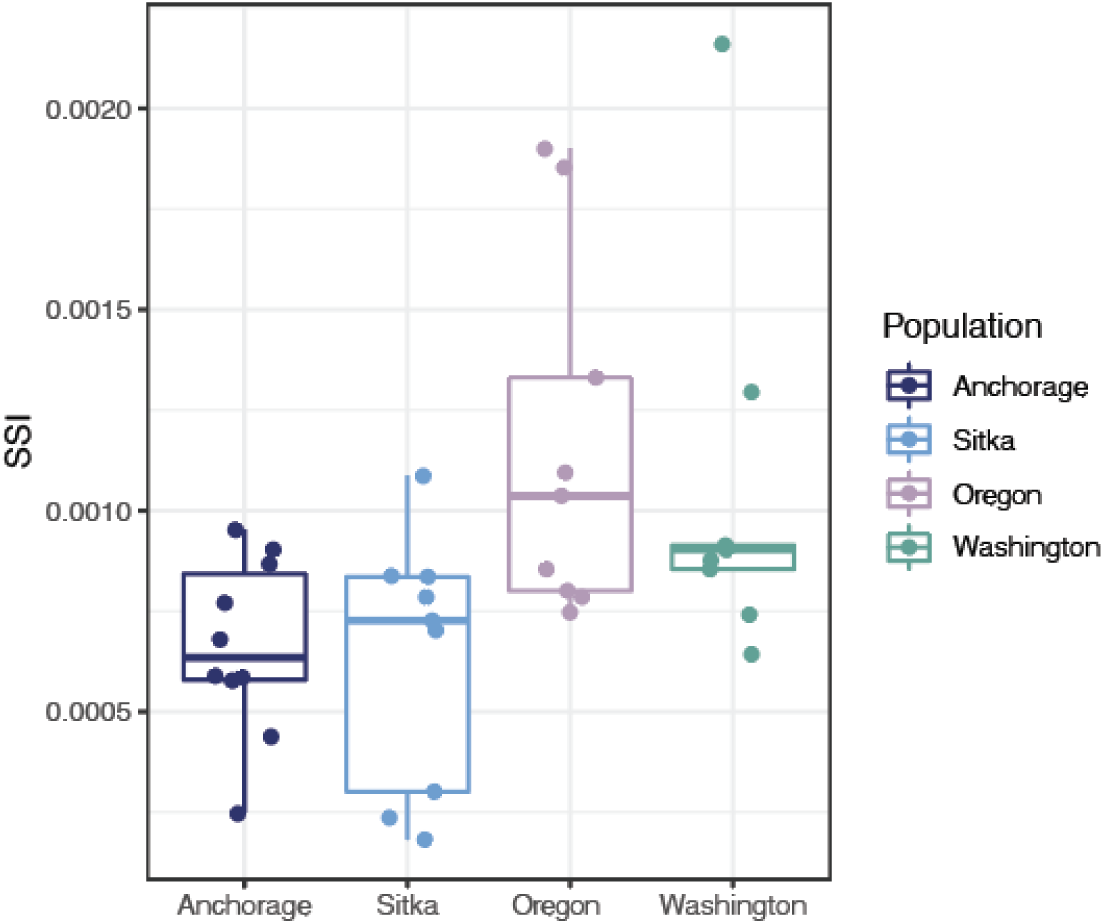
Splenoosomatic index (SSI) for each fish which underwent CT_max_ trials.

**Supplemental Figure 8:**
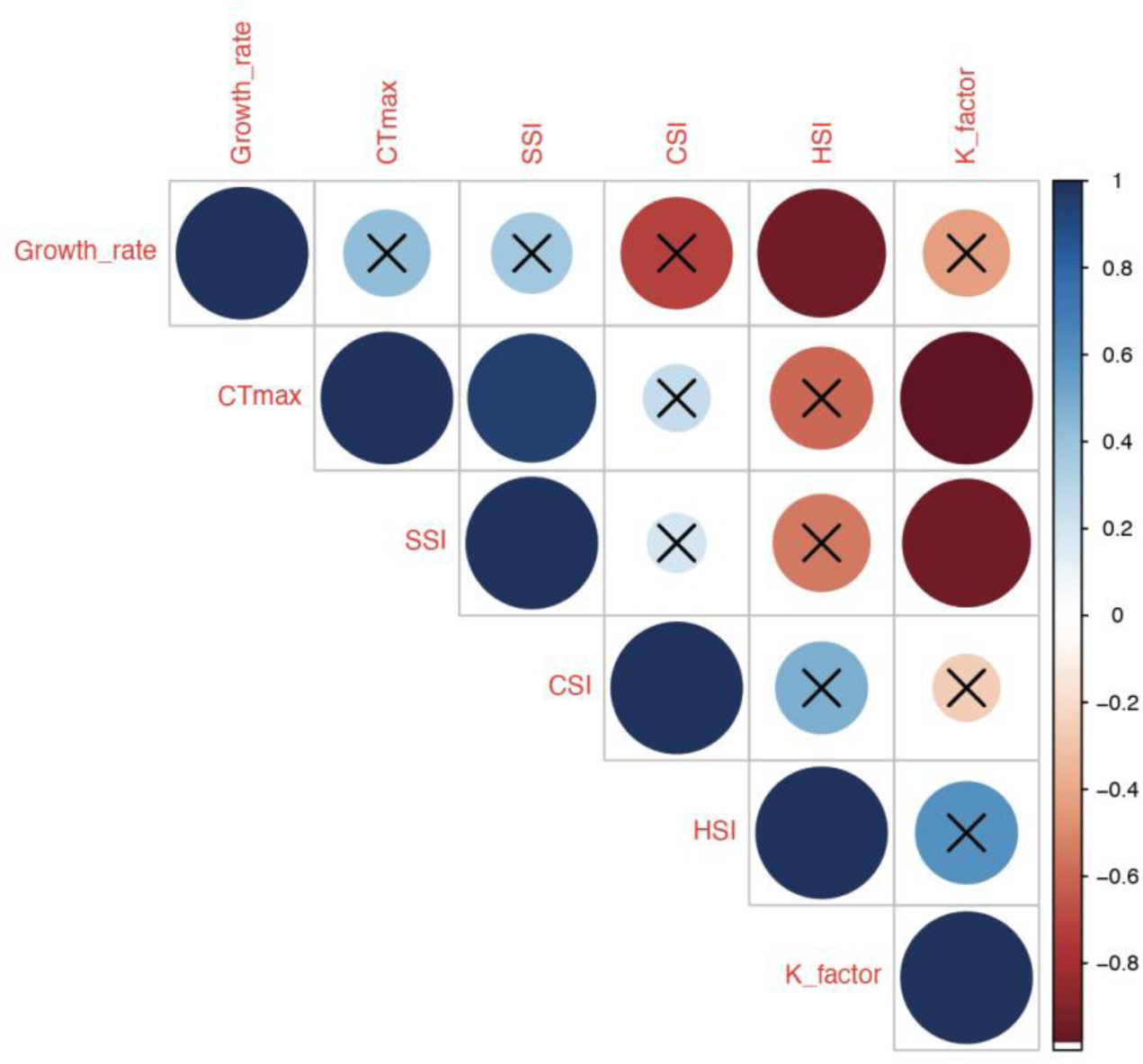
Correlation co-efficient between phenotypic measurements, where the strength of the correlation is depicted by both size and color of the circle. Non-significant correlations (p>0.05) have marks through them. Positive correlations are represented by blue colors where negative correlations are represented by red colors.

**Supplemental Figure 9:**
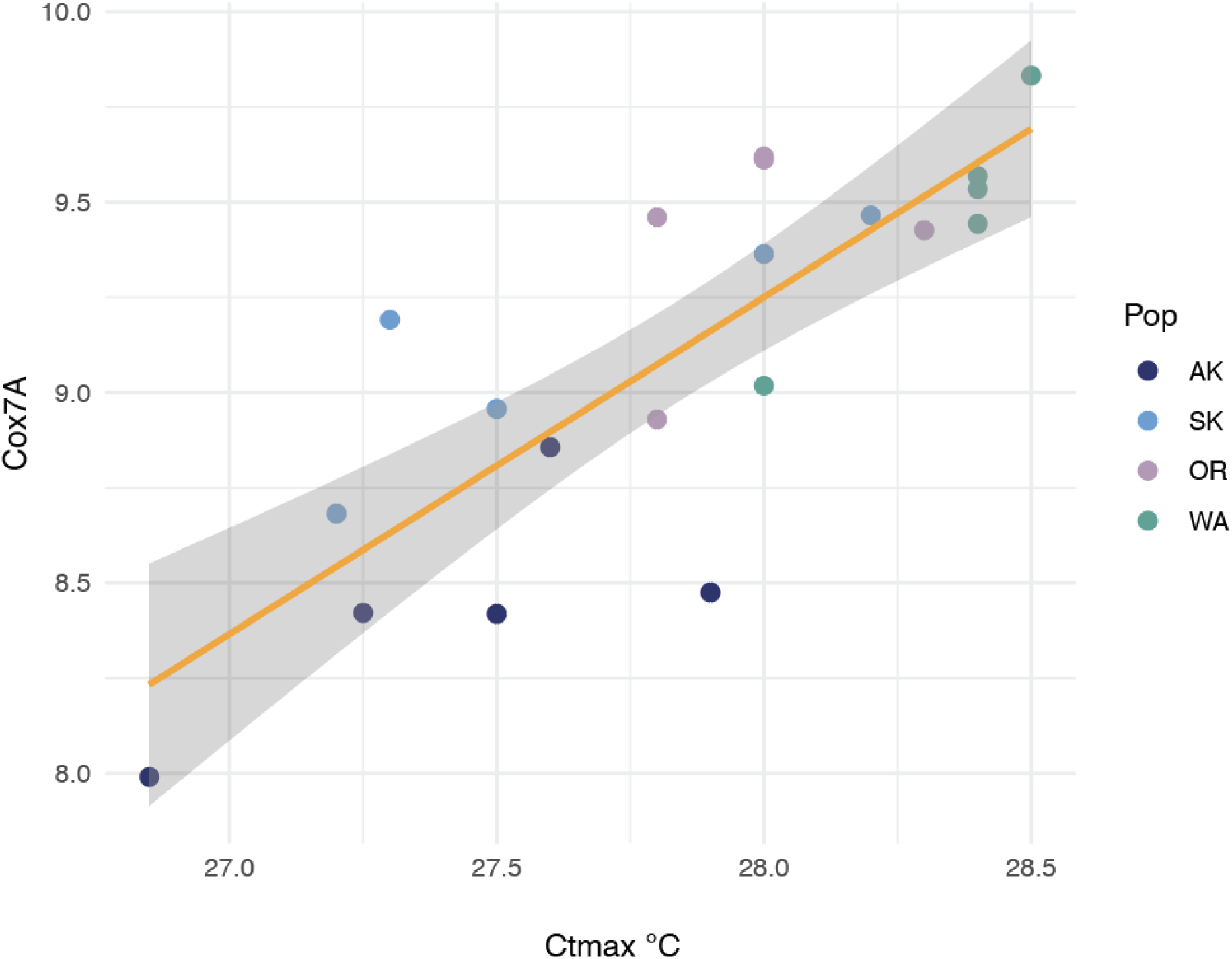
Correlation between the normalized expression of the Cox7a gene and fish CT_max_.

**Supplemental Figure 10:**
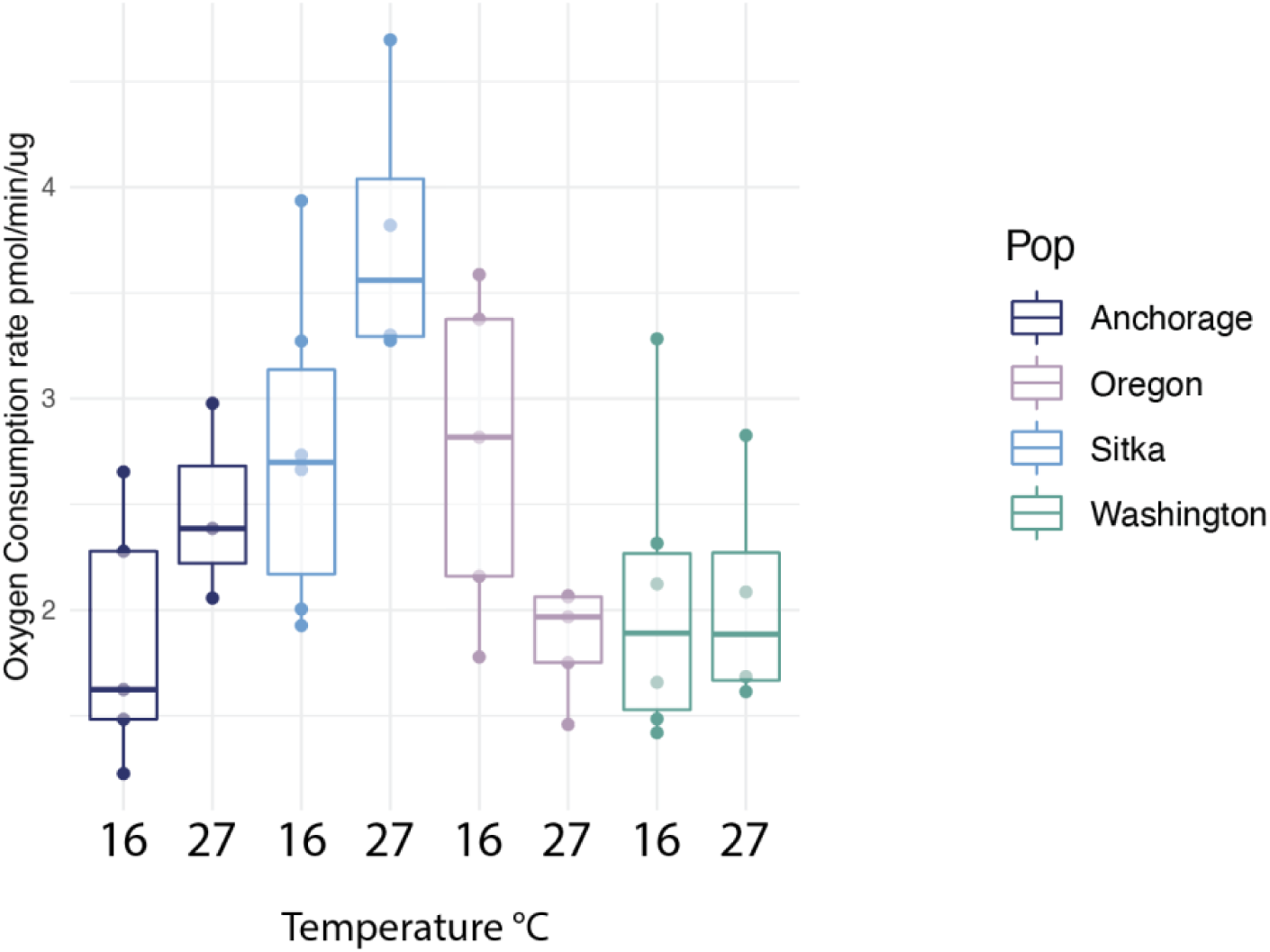
Basal oxygen consumption rate at 16 and 27.5°C for each population.

**Supplemental Figure 11:**
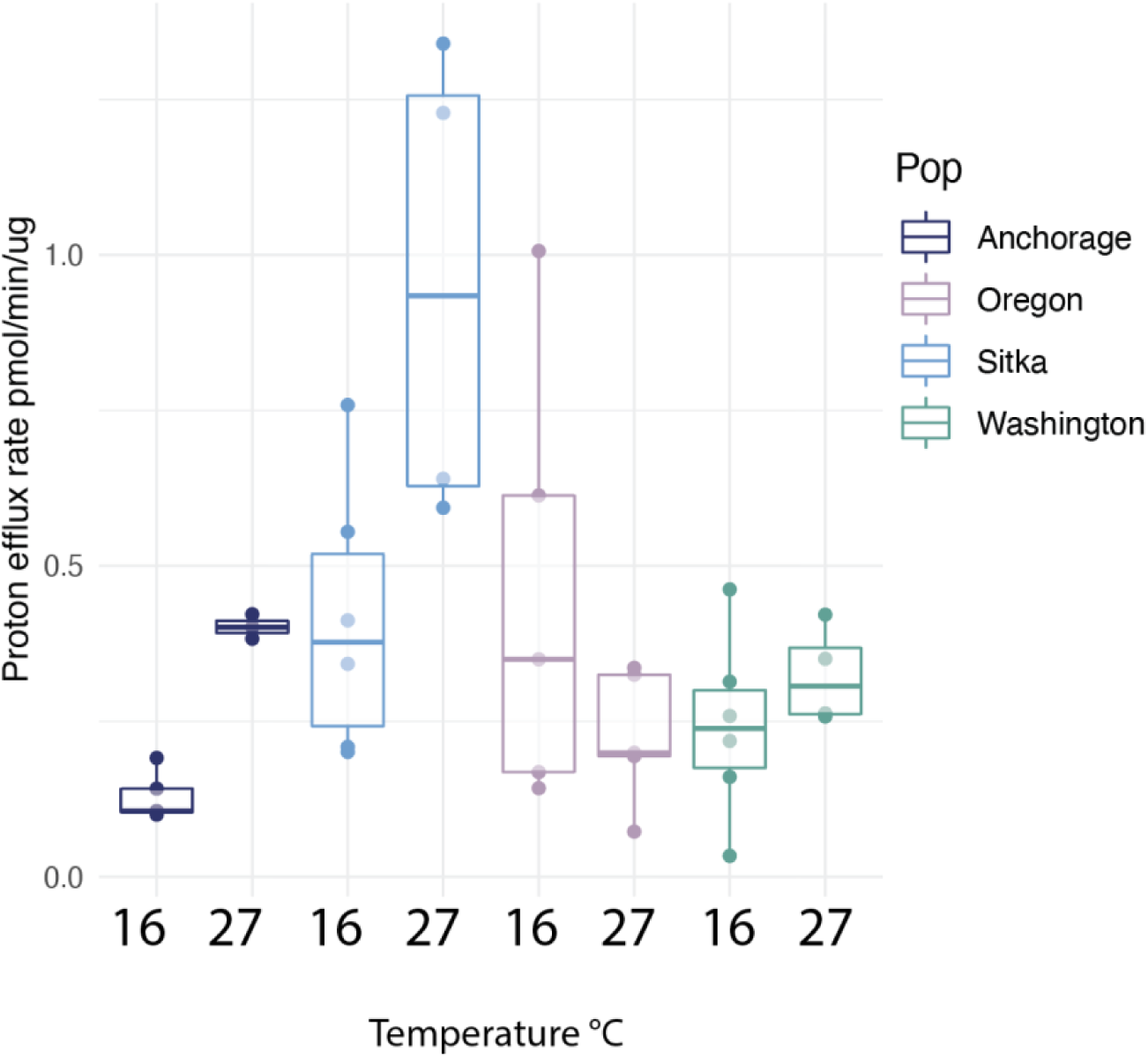
Proton efflux rate at 16 and 27.5°C for each population.

**Supplemental Figure 12:**
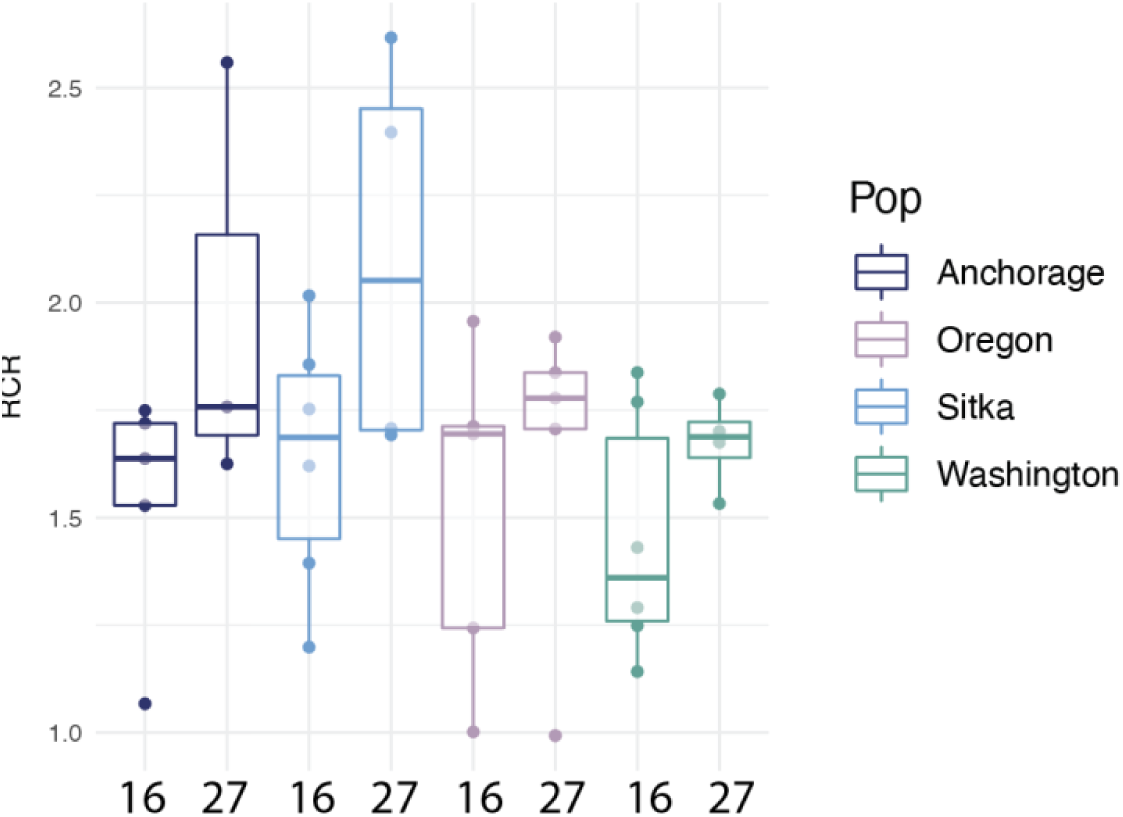
Respiratory Control Ratio (RCR) at 16 and 27.5°C for each population.

